# FINE GRAINED LONGITUDINAL ANALYSIS OF COCOA BEAN FERMENTATION PROVIDES INSIGHTS INTO THE DYNAMICS OF MICROBIAL POPULATIONS

**DOI:** 10.1101/702225

**Authors:** M. E. Pacheco-Montealegre, L. L. Dávila-Mora, L. M. Botero-Rute, A. Reyes, A. Caro-Quintero

**Author notes:** Address correspondence to Alejandro Caro-Quintero,.

## Abstract

Cocoa bean fermentation is an important microbial driven process where metabolites that affect chocolate quality and aroma are produced. Although considerable research has been devoted to the yeast and bacteria species involved in this process, less attention has been paid to the role of populations and strains, which hinders its selection, monitoring and use. Here we present a study that evaluates the microbial diversity associated to the tools and bean mass during spontaneous cocoa fermentation and in two distinct agro-ecological zones in Colombia. Using high-throughput sequencing of molecular markers for bacteria and yeast, we established the dynamics at the species-level (OTUs) and strains-level (oligotypes). Our results show that cocoa bean fermentation is catalyzed by a composite of strains within each OTU and not by one single strain. Eventhough we found only a few bacterial OTUs, one Enterobacter, three of Lactic Acid Bacteria and two of Acetic Acid Bacteria, these could be further split into 6, 23 and 19 oligotypes, respectively. Only two fungal OTUs were found. Comparison between fermentations suggest that local protocols generated a specific footprint in the dynamics of the microbial communities and that tools are reservoirs of some of those groups. The population analysis shows that the oligotypes that become most dominant are the same in the two locations, coupling co-abundance and dominance analysis we suggest that a combination of Enterobacter and Acetobacter oligotypes seem more optimal for the starter cultures. In conclusion, the results presented here show that exploring the fine level dynamics of microbial fermentation is necessary to understand the patterns of the dominance of specific populations and can be used as a valuable approach to select and monitor specific bacteria for the design of starter cultures in the food industry.

**IMPORTANCE:** In Colombia, the lack of tools to validate and standardize fermentation protocols are one of the principal reasons why Colombian cocoa beans are not recognized as “fine-flavor” and widely marketed. Because of the large influence of the microbial fermentation in cocoa quality, the present study explores the microbial dynamics using high-throughput sequencing of molecular markers in two of the most important producer regions in Colombia. The results show that the identification of dominant and transitive strains can be used to select, design and monitor starter cultures and/or the effect of adjustments in the fermentation protocols.

## INTRODUCTION

Cocoa bean fermentation is a spontaneous process where all bean structures (husk, mucilage, cotyledon, and embryo) are physically and biochemically transformed due to the activity of a microbial community composed by yeast, Enterobacteria (ENT group), Lactic Acid Bacteria (LAB group) and Acid Acetic Bacteria (AAB group) (1, 2). The fermentation process last three to seven days depending on variables such as the genetic origin of the seed, the agro-ecological conditions and the protocol used. Despite this variability, the fermentation process follows a predictable behavior. During the first hours, yeast degrades the sugars and pectin present in the cocoa pulp converting them into ethanol. Subsequently, LAB degrades the carbohydrate-rich pulp producing mainly lactic acid, finally, AAB transforms the lactic acid into acetic acid (3, 4). In this last step, the production of acetic acid lowers pH and increases bean mass temperature (40-50 °C), these conditions affect the seed and cause its death, promoting the subsequent release of the flavor precursors associated with chocolate quality (5–7). Microorganisms are also responsible for the production of volatile compounds related to aromatic flavor (e.g., alcohols, organic acids, esters, and aldehydes) and the removal of compounds associated with bitterness and astringency (e.g., like tannins, polyphenols) (1, 3, 8). The strong relationship between the ecological succession and the development of the desired organoleptic characteristics of chocolate makes cocoa fermentation an interesting system for the study of microbial ecology and a key process for the standardization and improvement of chocolate quality.

In the last decade, culture-independent methods have been incorporated into the analysis of the cocoa bean fermentation to track the relative abundance and the species composition of bacteria and yeast. In fact, techniques such as 16S rRNA gene PCR-DGGE (Denaturing gradient gel electrophoresis), 16S rRNA gene clone libraries, and PCR-RFLPs (Restriction Fragment Length Polymorphism) have brought new insights into the dynamics of the bacterial community involved in the fermentation process. Those analyses have shown a limited diversity in terms of the number of species involved in the cocoa bean fermentation (2), additionally, that most members of the bacterial groups present in the fermentation process are culturable, and that Enterobacteria are also consistently found in the process (9). Recently, metagenomic shotgun sequencing was used to evaluate the composition and metabolic potential of the community at one-time point of the fermentation (10), revealing a wider diversity of bacteria and yeast, and elucidating the contributions of the distinct functional groups in the biochemical transformation of cocoa bean mass. Despite the relevance of these studies, there are several shortcomings with the current methodologies, particularly, in the identification of population/strain level diversity, and the changes in their relative abundance over time throughout the fermentation process.

Identification of species and strains can be accomplished by the used of high-throughput sequencing of molecular markers. These methods allow a fine scale monitoring of compositional changes of the community tackling the limitations of previously used methods such as gel-based profiling and are a more cost-effective technique compared to shotgun metagenomics. In the case of high-throughput sequencing of 16S rRNA gene phylogenetic marker, species are defined as Operational Taxonomic Units (OTUs) (10), clustered at 97% sequence identity (11, 12). Identification of strains within each OTU is a more challenging task that cannot be achieved by identity thresholds, but by the use of newer computational analysis methods such as Oligotyping (13). This technique has been successfully implemented in a large variety of ecological studies such as, the human oral microbiota (14), plant-associated microbiota (15), the study of specific strains associated to infant food allergy (16), and simple microbiomes environments as Kombucha tea fermentation (17), among many more and have demonstrate its utility for the identification of meaningful sequence variants that provide ecologically significant subgroups. Recently, oligotyping analysis has been used to study the microbial community of fermented beverages (18), showing that it is possible to monitor closely related strains with different patterns of abundance and ecology.

Here we used high-throughput sequencing of phylogenetic markers to study the microbial communities associated to the cocoa bean fermentation process in two distinct agroecological zones (AEZs) from Colombia (19, 20), *Montaña Santandereana* (MS) and *Zona Marginal Baja Cafetera* (BC). Furthermore, we study the dynamics of the microbial community involved in the process of fermentation and the effect of protocols, the levels of inter-connectivity of the community with the tools employed during the fermentation process and the patterns of intra-specific diversity and its relationship with geographic location and/or fermentation protocols. Such understanding has important implications in the improvement of fermentation technologies, the optimization of protocols and might be relevant to monitor and select potential starter cultures (3, 4, 21).

## MATERIALS AND METHODS

### Sample Collection

From each AEZ, *Montaña Santandereana* (MS) and *Zona Marginal Baja Cafetera* (BC), one farm was selected for sampling. The criteria for selection was based on several factors including, the certification of Good Agricultural Practices (GAP), the existence of clear protocols of fermentation, and validation of fermentation through the monitoring of the structure and color of fermented beans using the cut-test (1) and determination of the fermentation index (**see Table S1 in the supplemental material**).

The sampling of the cocoa beans mass was done at two different times of the year (in 2016, rainy and dry season) at each AEZs (see **Fig. S1** in the supplemental material). Cocoa pods were harvest using brier hook tool or pruning shears, beans were removed by hand (Pulp-preconditioning phase) and placed in a fermentation wooden box and loaded with approximately 400 kg of cocoa beans. Other tools used in each farm are reported (**Table S2)**. In order to monitor the taxonomic composition of the microbial community, 100 gr of cocoa beans were collected from three different points at each of two different depths (upper and middle section) of the fermenter box using sterile gloves during the whole fermentation process (6-7 days). Samples were collected every 12 hours and immediately stored at −20°C. A total of 126 cocoa bean samples were collected.

A sampling of the microbial community associated with the harvest and postharvest process was done during the first sampling time to elucidate the transference and sink of the microbial community associated with the fermentation process. Sterile cotton swabs were used to collect the microorganisms associated with the surfaces of fruits, harvest tools and elements used during the fermentation process, after collection, each swab was immersed individually in sterile saline solution and stored at −20 °C until processing.

### DNA extraction

The microbial cells were recovered from the bean samples according to the protocol of Camu et al., (2), with some modifications. Twenty grams of frozen beans pulp samples in 250 ml Erlenmeyer flasks with 0.85% NaCl were homogenized in a rotatory shaker (150 rpm) for 30 min. The combined fluid was decanted and filtered through sterile gauze. The free-pulp solution was centrifuged at 3220 x *g* at 12 °C for 20 min to remove large particles. The biomass was resuspended in 10 ml of sterile distilled water. The DNA was extracted using the UltraClean® Microbial DNA Isolation kit and the concentration and purity of DNA was quantified using a NanoDropTM 1000. The final community DNA samples were stored at −20 °C until further use.

### Amplicon Sequencing

The analysis of the microbial community associated with the cocoa bean mass was done through amplicon sequencing of conserved molecular phylogenetic markers. For bacteria, the regions V3-V4 of 16S rRNA gene were amplified, while for yeast the region ITS2 of 5.8S – LSU operon was used. Bacterial and yeast DNA libraries were prepared according to Faith et al. (22). In brief, the sequencing library preparation was carried out in a two-step PCR procedure. In the first PCR, the modified primers 515F-806R (23) and ITS4_ KY03-ITS3_KY02 (24) were used for bacteria and yeast, respectively. Primer modification includes a linker region in the 5’ end of the primer, as shown in **Table S3**: During the second PCR barcodes are added to the amplicons as well as the illumina i5 and i7 regions (25).

The first PCR, for either gene or ITS region, was performed in triplicate and carried out in 25 µl reaction volumes. Each reaction contains 0,1 µl of Taq Platinum (Invitrogen), 0,75 µl MgS04 50mM, 2,5 µl Buffer-Mg 10X, 0,5 µl dNTPs 10mM, 0,5 µl (10µM) of each forward and reverse primers with adaptor sequences, 2 µl ADN and 18,15µl of Ultrapure Distilled Water (Invitrogen). In the case of the 16S rRNA, the first PCR starts with denaturation at 94°C for 120 s, followed by 35 cycles of denaturation at 94°C for 45 s, annealing at 50°C for 60 s and extension at 72°C for 90 s, and a final extension of 72°C for 10 mins. In the case of the ITS, the first PCR starts with denaturation a initial denaturation at 95°C for 120 s, followed by 35 cycles of denaturation at 95°C for 30 seconds, annealing at 55°C for 30 s and extension at 72°C for 60 s, and a final extension of 72°C for 5 mins. Both 16S rRNA and ITS Amplicons were visualized in agarose gels (1.5% p/v) in 1X TAE at 100 V, 30 mins. All PCR products were purified following the protocol given by Agentcourt® Ampure® XP

The second PCR use both amplicons from the first PCR as template and was carried out by adding 5 µl of purified product, 1 µl (10 µM) of each of the forward and reverse indexed primers, 8 µl of the 5-Prime HotMasterMix 2.5×, and 5 µl of UltraPure Distilled Water (Invitrogen). PCR was performed using the same reaction conditions as above but allowed to proceed for only 12 cycles. Products of the second PCR were purified with AMPure XP beads, and DNA concentrations were quantified on a Qubit 2.0 fluorometer (Invitrogen). Samples were normalized to equal concentrations, combined and pair-end sequenced (250-nt reads) on the Illumina MiSeq by the Microbial genomics laboratory of the Molecular Genetics and Antimicrobial Resistance Unit at Universidad El Bosque.

### Bioinformatics analysis

#### Pre-processing of reads

MiSeq sequencing reads were transferred to the HPC system at Universidad de los Andes. Quality inspection was performed using FastQC v. 0.11.2. The trimming was done with Trimmomatic v. 0.36, removing primers and nucleotides with low quality score. Demultiplexing and pair-end assembly were performed using QIIME v 1.9.1 scripts (join pair-end reads and split libraries) with high stringency in specific parameters (**Table S4**).

#### Operational Taxonomic Unit (OTU) determination

We used QIIME version 1.9.1 to perform *de novo* OTU clustering with UCLUST method (12) for both molecular markers. Subsequently, we perform taxonomic assignment against the Greengenes database (version 13.8, http://greengenes.lbl.gov), and created a biom table where we filter chloroplast and OTUs with abundance below 1%, for 16S rRNA gene reads (parameters are included in **Table S4**). For the ITS reads we used NCBI-BlastN against the nucleotide non-redundant (nt) database to perform the taxonomic assignment.

#### Oligotyping protocol for strain identification

Oligotyping v. 2.1 (http://merenlab.org/software/oligotyping/) was used for the generation of strain level groups, for this, all reads of an OTU were selected and aligned using PyNAST, as implemented in QIIME. Subsequently, we stripped common gaps from each alignment in accordance with the in-house pipeline (http://merenlab.org/2012/05/11/oligotyping-pipeline-explained/), and the Shannon entropy was calculated for each base position in the alignment (13). Finally, to resolve all oligotypes in a bacterial taxon, we used all available highly variable base positions for each OTU. For this purpose, we select specific Shannon-entropy cut-offs (**Table S5**) for each OTU and all positions in the alignment that present equal or better value were selected as components (c parameter) for the oligotype script in the pipeline.

An oligotype was considered if it occurred in more than 10% of all reads (a = 10) and if the minimum number of samples containing the oligotype was more than 10 (s = 10). The counts of each oligotype were normalized according to the total number of 16S rRNA gene reads per sample and for generating a heatmap visualization we used the package *pheatmap* in R, after a square root transformation of the reads counts.

To evaluate the relationship between all oligotypes from the study, we used the Spearman’s correlation coefficients (rs) with *Hmisc* and using *corrplot* for visualization, libraries in R.

#### Analysis of bacterial transference/connectivity

We used SourceTracker v. 1.0.1 (https://github.com/danknights/sourcetracker) to establish the possible sources of transference or inoculation (sinks) of bacteria associated with the fermentation process (26). For this, sequences from the first sampling time were obtained from one swab sample from each fermentation tool category (**Table S2**) and were used as sources and the sequences from the corresponding fermentation process were used as sink. All reads were used for *de novo* OTUs (UCLUST method) picking for both *BC* and *MS*. Subsequently, the generated biom table was filtered by minimum count fraction of 1% and any OTU assigned to chloroplast was removed. Following, sourcetracker_for_qiime.r was used following the developer’s suggestions (https://github.com/danknights/sourcetracker).

#### Data availability

The bacterial and fungal sequence data generated in this study using MiSeq have been deposited and are available in the NCBI Sequence Read Archive (SRA) under BioProject PRJNA492720.

## RESULTS

### Fermentation protocols varied between farms

Cocoa plantations in Colombia are distributed along the country and each productive region is delimited in agro-ecological zones based on climatic conditions, topography, and soil composition. We choose two of the most relevant AEZ, *Montaña Santandereana* (MS), the region with the largest rates production of cocoa in the country, and *Zona Marginal Cafetera Baja* (BC), one of the most productive regions with low temperature and humidity rates (20). For each AEZ one farm was selected for sampling. The criteria for selection was based on several factors as described in Methods and **Table S1**.

A detailed analysis of the fermentation protocols showed that the practices between farms are highly heterogeneous. The most notorious differences were the length of the process and the timing for initiation of the bean mixture. In MS, the length of the fermentation process was in general shorter 108 h in March (dry season), and 132 h in June (wet season), while in BC farm fermentation lasted 132 h in May (dry season) and 144 h in June (wet season) (**Fig. S1**). Mixing of cocoa beans also started at different times, 48h in May (dry season) and 24h in June (wet season) for BC while it started at 24h for both climate regimes in MS.

### Sampling and amplicon sequencing

Cocoa bean samples were collected throughout one complete fermentation process during two different climate regimes (wet and dry season). Sampling was done every 12 hours and at two depths in both farms. Overall, 94 samples were collected for the microbiota analysis, 44 samples from MS and 50 from BC farm.

Amplicon libraries for 16S rRNA gene and fungal ITS region were prepared and sequenced for each sample (Table 1). A total of 2,742,198 reads were obtained for 16S rRNA gene libraries. After quality control 1,426,240 reads were kept for further analysis. In the case of ITS sequencing 443,340 reads were obtained in total and after quality control 60,339 reads were kept. The low number of filtered reads for yeast and bacteria are a consequence of the astringency of the sequence quality control and trimming (**Table S4**).

**Table 1.** Summary of the sequencing effort during the microbial monitoring study of the cocoa bean fermentation

|  |  | Zona Marginal Baja Cafetera (BC) |  |  |  | Montaña Santandereana (MS) |  |  |  |
| --- | --- | --- | --- | --- | --- | --- | --- | --- | --- |
|  |  | Dry season |  | Wet season |  | Dry season |  | Wet season |  |
|  |  | May |  | June |  | March |  | June |  |
|  |  | Upper | Middle | Upper | Middle | Upper | Middle | Upper | Middle |
|  |  | zone | zone | zone | zone | zone | zone | zone | zone |
| BACTERIA | Number of raw sequences | 330,892 | 263,452 | 297,372 | 338,781 | 295,059 | 274,659 | 480,765 | 461,218 |
|  | Number of clean sequences | 242,225 | 185,386 | 224,372 | 249,150 | 214,897 | 213,455 | 345,905 | 314,520 |
|  | % of reads assigned to cocoa tree* | 3.21 | 5.06 | 4.14 | 4.18 | 7.92 | 0.99 | 5.02 | 7.11 |
|  | Number of Oligotypes | 44 | 45 | 47 | 44 | 42 | 48 | 48 | 48 |
| YEAST | Number of raw sequences | 83,678 | 64,715 | 66,344 | 62,764 | 56,342 | 32,586 | 34,624 | 42,287 |
|  | Number of clean sequences | 10,014 | 10,857 | 12,564 | 10,330 | 9,619 | 5,985 | 5,935 | 5,365 |

\* The value was calculated from the number of clean reads.

In order to determine the saturation of the sequencing effort, a rarefaction analysis was done for all individual libraries of 16S rRNA gene and ITS at the OTU level. The result shows that for 16S rRNA gene and ITS libraries, the diversity present in the samples is well covered (**Fig. S2**). A similar result was obtained for the ITS libraries (data not shown), showing that the obtained reads were also enough to saturate the yeast and mold diversity present in the samples.

### The fermentation process is governed by a low number of species

The bacterial taxonomic analysis identified six bacterial OTUs (97% identity) and one *Theobroma cacao* chloroplast OTU (in both farms). Around 5 % of the processed 16S rRNA gene reads were assigned to chloroplasts and were present mainly during the first 24 hours of the fermentation process (**Fig. S3**), these reads were removed from the microbial community analysis, but were used later to elucidate patterns associated to practices and protocols. During the initial hours of the fermentation the most abundant bacterial species corresponded to an OTU assigned to the *Enterobacteriaceae* family (Figure 1A and B). The Lactic Acid Bacteria became dominant mainly at the 24 to 36 hours of the fermentation, this group was represented by three distinct OTUs assigned to the genus *Lactobacillus sp*., the family *Lactobacillaceae*, and the genus *Fructobacillus sp*, respectively. Finally, the Acetic Acid bacteria, dominated after the 48 h in both farms, these group was represented by two OTUs assigned to the genus *Acetobacter sp* and the *Acetobacteraceae* family, respectively. Regarding the ITS assignments, four OTUs at the 95% identity-level were detected throughout the fermentation process (Figure 1C and D). *Hanseniaspora opuntiae* represents more than 80% of yeast relative abundance at any time during the fermentation process. *Pichia sp*, *Pichia kudriavzevii* and *Wickerhamomyces pijperi* raised slowly and appear only in the final stages of the fermentation process in both AEZs.

**Figure 1.**
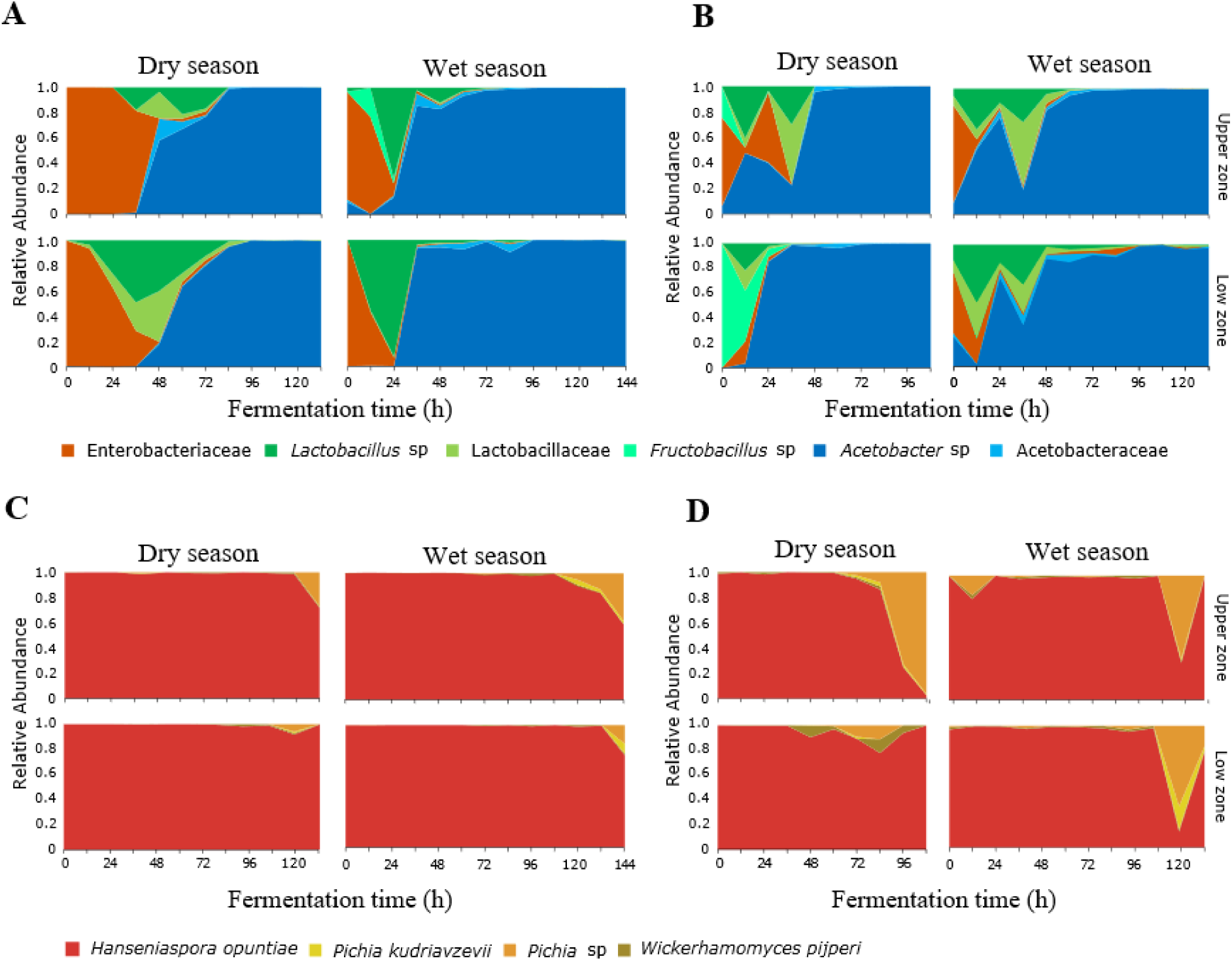
A predictable microbial succession of bacteria and yeast, during the fermentation process of cocoa beans. A total of 6 OTUs for bacteria (A, B) and 4 OTUs for yeast (C, D) were detected. Relative abundance for each OTU at the different Agroecological zones on each depth and month surveyed are shown during the first 140 hours of the fermentation process.

### Fermentation protocols affect the succession of the microbial community

The bacterial composition during the fermentation process showed differences between farms, especially in the time of transition and dominance of bacterial groups. In the BC farm during the dry season sampling (Figure 1A) the ENT group was present during the first 36 hours, being dominant during the initial 12 h, the LAB group was dominant at 48 h and the AAB group was the dominant OTU from the 56 h to the end of the process. While in the wet seasons sampling, the ENT was present only during the first 24 hours. LAB showed one clear peak of abundance at 24 h, while AAB was dominant after the 36 h. In the MS farm during the dry season sampling, the ENT was present during first 36 hours (Figure 1B), LAB were present during the initial 36 hours showing two peaks of dominance at 12 and 36 h. Finally, the AAB were dominant after 48 hours and until the end of the process. In June, the ENT was present during the first 12 hours, the LAB were present from 0 to 48 hours, and similarly to what happened in March, two peaks of abundance were observed. Finally, the AAB were dominant from the 48 hours to the end of the process.

The variation in the times of transition in the microbial succession seems to be affected directly by the fermentation protocols used. For instance, the two farms start bean mixing (the incorporation of oxygen) at different periods of time (e.g. 24 h in MS and 48 h in BC) which affects the transition of LAB to AAB group. This is clearly shown by the effect of bean mixture in the increase of temperature and AAB abundance (**Fig. S4**). Other fermentation practice that affected the dynamics of the microbial community is the posterior addition of fresh bean mass to an ongoing fermentation process. This practice is evident in the MS fermentation during both seasons, where the addition of freshly harvested seeds can be detected by the presence of *T. cacao* chloroplast in the 16S rRNA gene amplicon libraries. Under regular fermentation conditions the cocoa seed dies and the presence of the chloroplast OTU decreases in abundance compared to the bacterial OTUs (**Fig. S3B and C**), however in the MS fermentation, chloroplast abundance increases at 36 hours, accompanied by a second unexpected peaks of abundance of LAB which coincides with the idea of the addition of fresh material by the farmer.

### The microbial composition involved in the fermentation is not homogenous at the surface and middle zone of the bean mass

The microbial composition was compared between the upper (surface) and middle zone (internal bean mass) to explore how the exposition to external conditions affects the dynamics of the microbial community within the same fermentation process. The comparison was done by subtracting the middle zone relative abundance of each OTU from their abundance in the upper zone; values below zero represented higher abundance in the middle zone, while values greater than zero indicated higher relative abundance in the upper zone.

The comparison between the relative abundance of upper and middle zone shows heterogeneity in the dynamics of the microbial community in both farms (**Fig. S5**). The relative abundance analysis shows that LAB group appears faster and becomes more abundant in the middle zone than in the upper zone independently of the seasons and farm, in particular during the first 36 hours. In contrast, the ENT group has a higher abundance in the upper zone during the first 12 to 14 hrs. These results also show that transition from LAB to AAB is faster in the upper layer. These patterns do not seem to be the result of differences in temperature as the analysis of temperature profiles between zones show no significant difference (**Fig. S4**), which suggest that the different environmental exposition (e.g., higher oxygen concentration) may be responsible for the heterogeneity in the microbial succession and might be responsible for the variability on bean fermentation.

### Higher resolution of bacterial oligotypes identifies dominant variants

The oligotyping analysis was used to detect informative variants (inter-specific diversity) for each bacterial OTU (13) and to establish the intra-specific variation between locations, sampling seasons and along the fermentation process. The use of this method allowed the detection of 48 oligotypes in total (Table 1). A total of 30 oligotypes were identified for OTUs with taxonomic assignment at the family level: 12 for the *Lactobacillaceae* OTU, 12 for *Acetobacteraceae* OTU, and 6 for *Enterobacteriaceae* OTU. In the case of OTUs assigned to genus, 18 oligotypes were found, 7 assigned to the *Lactobacillus* sp OTU, 7 to the *Acetobacter* sp OTU, and 4 to the *Fructobacillus* sp OTU (Figure 2 **and Table S5**), showing that higher taxonomical assignment does correlate with a higher intra-OTU diversity In general, the higher diversity of oligotypes was observed at the beginning of the transition between functional bacterial groups (ENT to LAB, and LAB to AAB). The highest values of Shannon diversity were observed after 24 hours where LAB were the most abundant group and the lowest was observed after 72 h where a few AAB oligotypes were dominant (**Fig. S6 and Table S5**).

**Figure 2.**
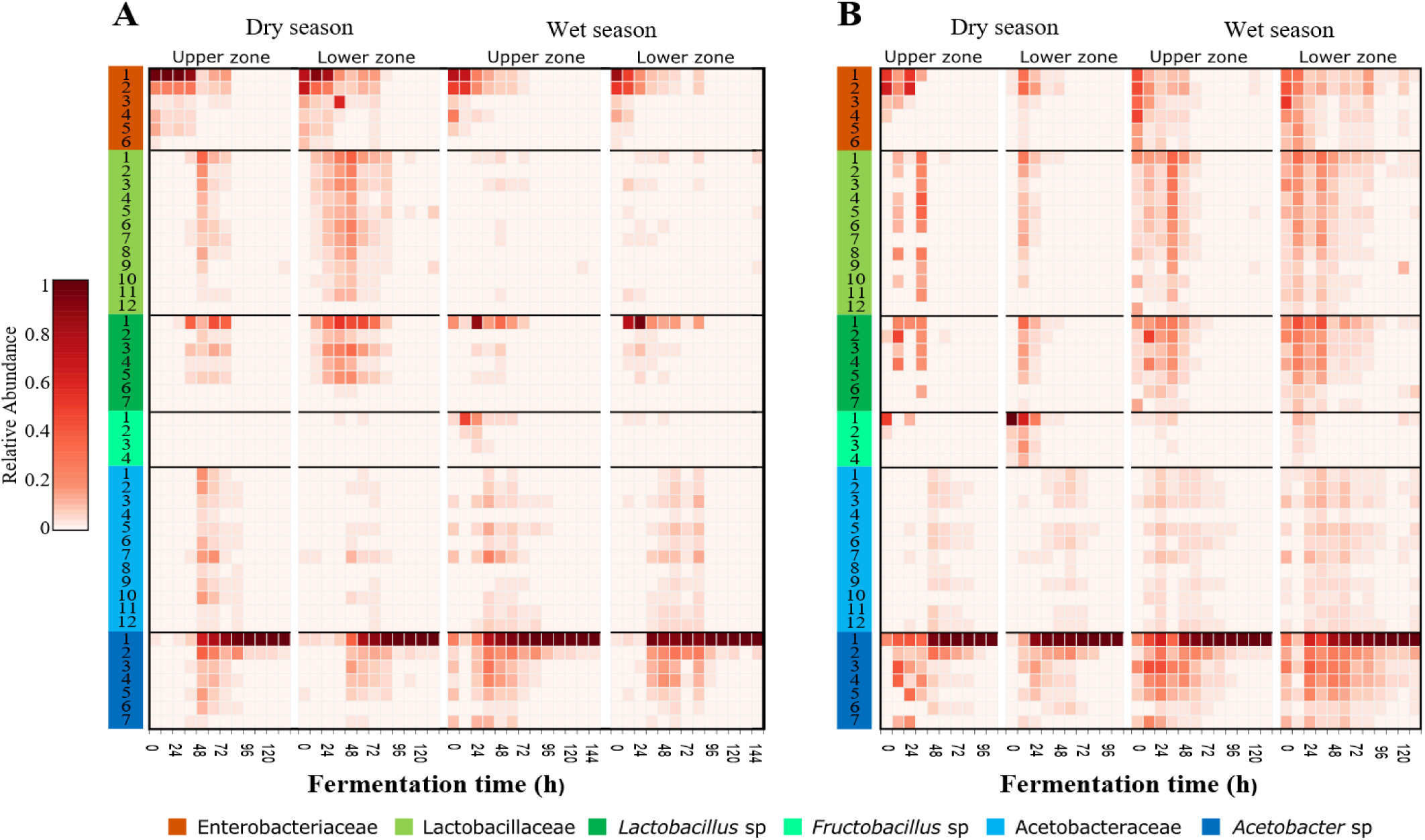
Dominant and transient strains of bacteria in each process of cocoa bean fermentation of both agroecological zones. Heatmap of the log transformed relative abundance for the identified oligotypes (strains) for each fermentation process of both agroecological zones (A, B) at each month and depth surveyed. Strains (rows) abundance is represented as a function of time (columns). Each oligotype observed is colored at the leftmost column according to the original OTU taxonomical assignment and the number corresponds to the Table S5.

The specific analysis of oligotypes diversity and abundance within the different groups, ENT, LAB and AAB allowed us to identify dominant and secondary oligotypes. In general, the most abundant oligotypes of each group were present in both farms and both sampling seasons (Figure 2). In the case of the ENT group, the patterns of abundance and the level of dominance changed between farms, where the oligotype Enterobacteriaceae-1 was more abundant and dominant in BC, being 2 to 13 times more abundant than the Enterobacteriaceae-2 oligotype. In contrast, in MS there is higher evenness between the ENT oligotypes, where the most abundant oligotype throughout fermentation was Enterobacteriaceae-2 oligotype being 1.2 to 1.4 times more abundant than Enterobacteriaceae-1. In the case of the LAB group there was a higher variability of the most abundant variants in MS than in BC. In the former, the most abundant oligotype was Lactobacillus-1 being 3 to 147 (**table S5**) times more abundant than the second most abundant oligotype from the same group. In MS, Fructobacillus-1 was the most abundant LAB oligotype during May sampling, while in June there was a more even distribution of oligotypes and the most abundant one differed between upper and middle zones, being Lactobacillus-1 more abundant in the upper zone and Lactobacillus-2 the most abundant in the middle zone. For the AAB group, the same oligotype, Acetobacter-1, was found to be dominant for both farms and sampling times, being 24 to 127 times more abundant than the second most abundant oligotype from the same group.

### The patterns of diversity, dominance and interaction for starter culture selection

The inference of ecological interaction between oligotypes can be an informative criterion to select a starter culture and predict the outcome of the addition of the isolates to a spontaneous cocoa fermentation system. Such interactions can be inferred using the relative abundances of the different populations through times and locations (23). Given that most oligotypes were present in both AEZ and that a future starter culture should be functional in different regions, we analyzed the abundance correlation (coexistence) between oligotypes in both AEZ and used this correlation matrix as a proxy to determine the positive, neutral and negative correlations between the oligotypes (Figure 3).

**Figure 3.**
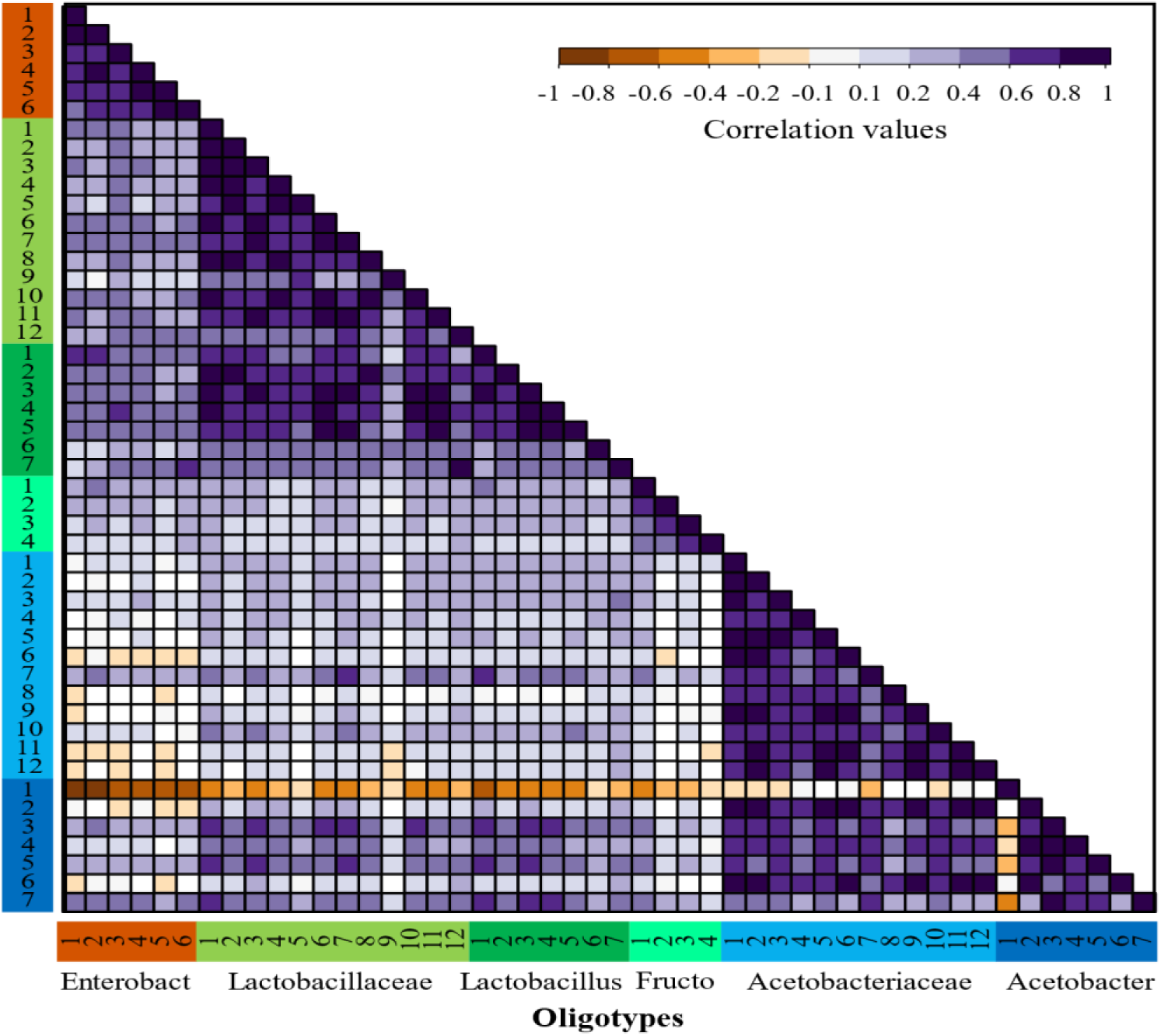
Correlations between strains of bacteria OTUs considering the oligotypes abundances in all fermentations process. Spearman correlations matrix between all bacteria strains detected in both agroecological zones datasets. Oligotypes are sorted in the same order and colored by OTU as shown in Figure 2 and the number corresponds to the Table S5.

The patterns of correlation reflecting possible positive interactions were observed more often among oligotypes obtained within similar functional groups (ENT, LAB and AAB), however, this was not the case for all the functional groups. For instance, for the ENT there is a strong positive correlation between Enterobacteriaceae-1 and Enterobacteriaceae-2, which is consistent in all farms. Within the LAB functional group, the Lactobacillus and Lactobacillaceae oligotypes have a significant positive correlation. In contrast, the low correlation was observed between these two former groups and the *Fructobacillus* oligotypes. In the case of AAB, the Acetobacteraceae oligotypes show a strong positive correlation with most *Acetobacter sp* oligotypes, nonetheless, there is a significant negative interaction between the most abundant oligotype from the *Acetobacter sp*, Acetobacter-1, with almost every other oligotype. This negative correlation can be observed in Figure 2 by its dominance in all fermentation processes at the last time points, where usually no other oligotype is observed.

The dominance indexes were also evaluated within each the bacterial group (AAB, ENT and LAB) to establish which groups might be better candidates for the isolation of starter cultures. We assumed that groups with high dominance indexes might be better for the isolation of highly abundant oligotypes and show a higher potential of success as we assured that the isolate is highly competitive. To do this, we quantified the distribution of the dominance index within each group. The samples used for this quantification were the ones where the relative abundance of the group was higher than 10 % to avoid stochastic effects (**Fig. S7**), The figure shows high dominance distributions for AAB with a very skew distribution where the median of the dominance index is higher than 0.9, which agrees with the patterns of correlation observed for the Acetobacter-1 oligotype (see above). In the case of the ENT group the distribution of the dominance index seems to be more intermediate with a median higher than 0.5, but with a bimodal distribution of the data with higher density around the dominance index of 0.5 and 0.8, this seems to be the results of the intermittent dominance of the Enterobacteriaceae-1 and Enterobacteriaceae-2 oligotypes. In the case of LAB, the violin plot shows a distribution skew towards values of low dominance with a median lower than 0.3, which agrees with the high diversity observed for this functional groups and the lack of a clear dominant oligotype within the group.

### Clustering fermentation processes by geographic location using correlation matrices

Correlation matrix between oligotypes using the relative abundances of all samples were used to compared fermentations as this approach allowed us to control the effect of different speeds of fermentation and focus on the interaction of the bacterial community. In total eight matrices were generated (as we had two farms, two seasons and two depths). The correlation between matrices was quantified using a Mantel test (27) implemented in PAST (28), the values of correlation were used to cluster the fermentation and to calculate the Bray Curtis index and a visualization through Principal Coordinate Analysis. The multivariate analyses of the correlation between all fermentation processes show a clear clustering by farm and season (**Fig. S8**). The Principal Coordinate Analysis shows an 81% variation explained within the first two axes, 56 and 25 % of the variability, respectively. An ordination plot of these axis shows that the first axis splits fermentation per farm, while the first and second axis cluster fermentations by time. Therefore, the analysis shows the clustering of fermentations from the same farms and season.

### Tracking bacteria colonization in the cocoa bean fermentation process

Identifying the potential source or origin for the different microorganisms that make part of the fermentation process could identify important contamination or inoculation points for the process. In this case, however, it was not possible to sample all possible tools, surfaces, and objects that were in contact with the beans during the fermentation process, only a representative tool from the harvest, pulp-preconditioning and fermentation process were used as potential sources of inoculation (**Table S2**). SourceTracker was used to estimate the potential origin for each of the dominant bacteria that take part during the fermentation process (Figure 4). The category “unknown origin” is obtained when the source tools do not fully explain the origin of a given taxon. This was particularly common in the mid-section of the fermentation, in particular when the LAB were dominant. The lack of a significant prediction of origin for LAB bacteria is likely due to the absence of such bacteria from the evaluated tools (**Fig. S9**). On early stages of the fermentation, when the Enteric bacteria are dominant, the main transfer appears to be with the harvest tools, in contrast, towards the end of the fermentation process when the AAB dominate, they seem to be transferred between the pulp-preconditioning tools and the fermentation boxes.

**Figure 4.**
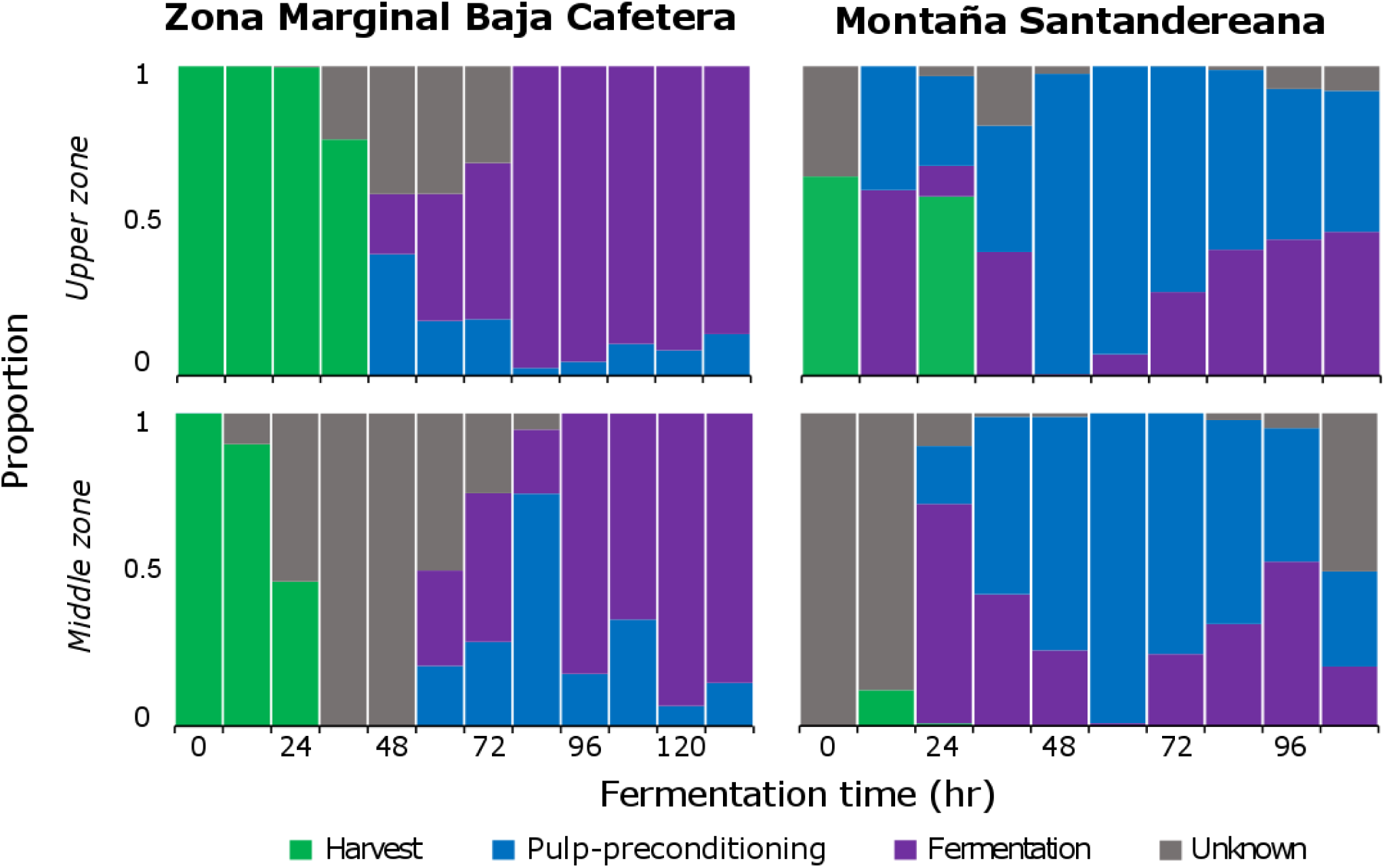
An *in-silico* prediction of the potential sources of bacteria during the first sampling months for both agroecological zones. SourceTracker was used to measure the probability of origin for the different bacterial species identified at each time point during the fermentation process at all AEZ for the first season at both depths and track them to either of 3 potential sources. For each sampling time, a relative proportion of origin is assigned between the 3 potential sources or an unknown source.

## DISCUSSION

### Microbial community analyses in the fermentation process for both agroecological zones

This study examined in detailed, in terms of sampling times and taxonomic resolution, the bacterial and yeast species populations present in two independent AEZ at two different seasons in Colombia. The bacterial and yeast species found in both farms corresponded to those previously reported in fermentation process from Brazil (29, 30), Ecuador (31), Ghana (9), Indonesia (32), Malaysia (33) and Mexico (34). Furthermore, the microbial composition and succession associated to the cocoa fermentation is in agreement with the reports obtained with methodologies such as PCR-DGGE and RFLP (34, 35), in particular, identifying bacterial and yeast groups such as Enterobacteriaceae, Lactobacillaceae, *Lactobacillus* sp, *Fructobacillus* sp, Acetobacteraceae and *Acetobacter* sp., *Hanseniaspora opuntiae*, *Picha* sp., *Pichia kudriavzevi* and *Wickerhamomyces pijperi*.

The presence of the Enterobacteriaceae group early in the fermentation has been reported in Ivory Coast (8) and Brazil (29). Additionally, Hamdouche et al., (8)and Papalexandratou, et al., (29)saw an association of the presence of Enterobacteria with Lactic Acid Bacteria members during the first 48 hours, similar to what we observed in almost all the fermentation processes. Enterobacterial species may contribute to glucose conversion into lactic acid and citric acid, and pectinolytic degradation of cocoa pulp (8), thus providing important resources for the Lactic Acid bacteria, furthermore, LAB metabolism relies exclusively on an anaerobic environment (6), which is likely caused after the fermentation process of ENT bacteria and yeast, explaining its rise following the Enterobacteriaceae expansion. The Acetic Acid Bacteria group dominated starting from 48 and 24 hours for BC and MS, respectively (Figure 1). The presence of this group in the last fermentation hours is similar to other reports(5, 7, 36), transforming the lactic acid into acetic acid, lowering pH and increasing bean mass temperature (40-50 °C), this last process is promoted by the mixing of the material by the farmer.

The use of deep sequencing of genetic markers to study the microbiota associated to the fermentation not only allowed to identify the species, but also provides a tool to monitor the process, for instance, the transition between groups, the unwanted practices by the farmers and the heterogeneity inherent to an open fermenter with uncontrolled conditions. In this sense, farmers decision of when to start the bean mixing alters the succession between LAB and AAB, the production of acetic acid and the increase the in temperature in the fermenter. Another identified practice (protocols) that affects the microbial composition is the addition of fresh collected beans into an ongoing fermentation process (**Fig. S3**). Evidence of this practice was observed at MS, by the detection of 16S rRNA genes from the Chloroplast at the 36 and 48 hours after the expected decreased in the Chloroplast reads at time 12 hr. The increase on Chloroplast abundance was accompanied by the decrease in AAB and increase of LAB, opposite to the expected succession (6, 37), and likely an indicator of an altered fermentation process.

The comparison of microbial communities between the upper and middle zone of the cocoa bean mass within the same fermentation showed a variation on LAB abundance and the transition times. The lower abundance of LAB in the upper zone is likely the consequence of its low oxygen tolerance (38), that prevents efficient colonization of LAB in the most external cocoa bean layers. Which in turn will modify the bean mass transformation rates. This heterogeneity is expected, because fermenters used by the farmers in most parts of the world are open systems where environmental conditions are harder to controlled.

### Strain community analyses in the fermentation process for both agroecological zones

The OTU level diversity within and between the cocoa fermentation processes from both agroecological zones agrees with the low diversity described previously in other studies (Figure 1), however, a closer look at the microbial diversity using the Minimum Entropy Decomposition (MED) (39)as implemented in the oligotypes approach, allowed us to examine with higher resolution how stable fermentation processes affect the intra-specific diversity of dominant bacterial groups in: different times, zones of the box, time of the year and agroecological zone (**Table S5**).

In contrast to the low OTU diversity, we observed high intra-specific diversity (Figure 2) with characteristic patterns of variation as a function of time. The dominant oligotypes for each group were the same in the two farms. Diversity seems to be higher at the beginning of microbial transitions in the succession, in particular for the new established functional groups, and decreases over time with few dominant oligotypes being selected. Such behavior can be clearly observed with the AAB group, when evaluating the dominance and Shannon indexes throughout the fermentation process (**Fig. S6**) suggesting that at the beginning of the succession there are more niches to colonize by individual strains but as the fermentation process continues, conditions change reducing the intraspecific diversity. Such results can be driven by physicochemical changes (e.g. pH, temperature), nutritional changes (depletion of nutrients), or by direct competition with the dominant OTUs. The dominance patterns suggest that in order to design starter cultures, only a few isolates of each group might be necessary; however, experiments are needed to elucidate the importance of non-dominant bacteria in the fermentation and the generation of chemical precursors of cocoa quality.

Although interaction and competition might be happening between oligotypes within groups (ENT, LAB and AAB), almost no interaction is evidenced between groups. Only one clear negative interaction was observed for AAB oligotype Acetobacter-1 and the other strains and is more likely a consequence of the combined effect of environmental conditions that exclude other functional groups such as ENT and LAB and competition or microenvironment conditions that removes other AAB oligotypes. Within the LAB group, *Lactobacillus* and *Fructobacillus* have little interaction or competitive exclusion based on abundance correlation, the differences of dominance and relative abundance in sympatry might reflect resource partitioning and micro-niche adaptation (11).

A better understanding of the patterns of diversity within species, is necessary to establish the relationship between strain diversity and metabolic function. In fact, there is evidence that suggests that not all strains have the same role and vary in their metabolic contribution to the fermentation process. For instance, it was identified through metafluxome analysis that *Lactobacillus fermentum* strain NCC575 uses fructose as an alternative external electron acceptor while *L. fermentum* NCC52 do not (40) and therefore there is a clear difference in their abundance and activity during cocoa fermentation.

Finally, fermentation processes are not entirely homogenous across regions, in some cases the transition of the ecological successions happened faster, and in consequence some fermentation ended earlier than others, these differences difficult a direct time-point to time-point comparison between locations. Here we show that co-abundance matrices between oligotypes can be used to solve this issue by comparing instead the interaction between the bacterial populations. This showed a clear clustering of samples by the farm, reflecting either the local conditions effect (e.g., temperature, humidity) or the effect of the different fermentation protocols used by the farmers. In any case, microbial interactions derived for patterns of oligotype co-abundance seem to be specific and therefore reflecting a strong relationship with location.

### Bacteria colonization in the cocoa bean

Identification of the source of the microorganisms in a given community through the use of SourceTracker is a valuable tool but is dependent on an exhaustive exploration of potential origins. In our case, even though we had a low number of observed species, it was not possible to predict the origin of all of them. In particular, the LAB group was not detected from the tools sampled. Furthermore, the taxonomical assignment of the tools samples shows that Enterobacteriaceae are highly abundant on harvest related tools for both zones sampled (**Fig. S9**) in agreement to reported members of this family, *Tatumella* sp and *Pantoea* sp, that are bacteria associated to fruits of the cocoa tree (41, 42).

For the origin of the AAB, dominant at later stages of the fermentation, there is some uncertainty on the results, since bacteria from the AAB group were identified in both the fermentation and Pulp-preconditioning sampled tools. Hence the mixed results obtained by SourceTracker for the final segment of the fermentation process. A further effort in characterizing other tools and components related to the fermentation process may thus be necessary to accurately predict the source of colonizing microorganisms.

## CONCLUSIONS

This is the first study on the microbiome dynamics of cacao fermentation that is performed at high level of resolution in terms of times (every 12 hours for up to 10 days) and taxonomic resolution, using deep marker gene (16S rRNA gene and ITS) sequencing for oligotype analysis. Two main contrasting observations were observed, first, at coarse level, the fermentation process is extremely conserved with the same functional groups, and even dominant oligotypes, being detected in all the different fermentations analyzed, with predictable successions over time. Second, at fine level, heterogeneity of the process is observed in terms of the exact time of microbial transitions and oligotype dominance and diversity, dependent on specific environmental conditions such as oxygen availability and addition of fresh material. Given the results, it suggests that following studies must analyze the microbiota associated with the fermentation at high resolution since those subtle changes may have a significant impact in the final product.

Microbial ecology studies in the food industry have the potential to guide better decision making towards the selection of microbial consortia for specific tasks. In our case, the dominance and co-abundance matrices allowed us to identified dominant populations within OTUs, for selection as starter cultures and for monitoring the fermentation process. The strains detection can be implemented to select, monitor and validate inoculation of specific strains to modulate and improve chocolate quality. Some of the microorganisms we observed in our study have been suggest as microbial starters (21, 33, 43), such as, *Acetobacter pasteurianus* and *Lactobacillus fermentum* (7, 33), additionally, some of the yeast groups (*Hanseniaspora opuntiae* and *Picha* sp.) have been reported in the fermentation process from other countries (6) and some have been used as culture starter (44). Underlining the importance of these microbial species for a successful process and their potential use as a microbial starter. Future perspectives, will direct these studies towards the design of family-specific molecular markers, e.g. PhyloTAGS(25), that provide higher resolution than 16S rRNA gene allowing monitoring of starter cultures in controlled and spontaneous conditions, and to understand the mechanisms for higher dominance of specific populations.

## ACKNOWLEDGMENTS

This study was supported by the agriculture and rural development Ministry of Colombia. We are grateful to AGROSAVIA and the HPC cluster from Universidad de los Andes, for invaluable help with the sampling and laboratory methodology and for computing support, respectively. The Max Planck Tandem Group in Computational Biology covered MEPM tuition during his master’s studies.

## CONFLICT OF INTEREST

The authors declare not conflict of interest.

## AUTHORS CONTRIBUTIONS

M.E. Pacheco-Montealegre performed all the data and computational analyses and statistics, discussed the results and wrote the manuscript. L. L. Dávila-Mora and L. M. Botero-Rute performed the sample collection and amplicon metagenomic sequencing. A. Reyes mentored on bioinformatics analysis, participated on the discussion and writing of the manuscript. A. Caro-Quintero did the experimental designed, participated in the sample collection, and advised on bioinformatic analyses, results discussion and writing of the manuscript.

## Notes

Conflict of Interest: The authors declare no conflict of interest.

## REFERENCES

1. Okiyama DCG, Navarro SLB, Rodrigues CEC. 2017. Cocoa shell and its compounds: Applications in the food industry. Trends Food Sci Technol 63:103–112.

2. Camu N, De Winter T, Verbrugghe K, Cleenwerck I, Vandamme P, Takrama JS, Vancanneyt M, De Vuyst L. 2007. Dynamics and biodiversity of populations of lactic acid bacteria and acetic acid bacteria involved in spontaneous heap fermentation of cocoa beans in Ghana. Appl Environ Microbiol 73:1809–1824.

3. Kongor JE, Hinneh M, de Walle D Van, Afoakwa EO, Boeckx P, Dewettinck K. 2016. Factors influencing quality variation in cocoa (*Theobroma cacao*) bean flavour profile – A review. Food Res Int 82:44–52.

4. Ozturk G, Young GM. 2017. Food Evolution: The Impact of Society and Science on the Fermentation of Cocoa Beans. Compr Rev Food Sci Food Saf 16:431–455.

5. De Vuyst L, Weckx S. 2016. The cocoa bean fermentation process: from ecosystem analysis to starter culture development. J Appl Microbiol 121:5–17.

6. Schwan RF, Pereira GV de M, Fleet GH. 2014. Microbial Activities during Cocoa Fermentation, p. 129–192. *In* Schwan RF, Fleet GH (eds.), Cocoa and coffee fermentations, 1st ed. CRC Press, Londres, UK.

7. Illeghems K, Pelicaen R, De Vuyst L, Weckx S. 2016. Assessment of the contribution of cocoa-derived strains of *Acetobacter ghanensis* and *Acetobacter senegalensis* to the cocoa bean fermentation process through a genomic approach. Food Microbiol 58:68–78.

8. Hamdouche Y, Guehi T, Durand N, Kedjebo KBD, Montet D, Meile JC. 2015. Dynamics of microbial ecology during cocoa fermentation and drying: Towards the identification of molecular markers. Food Control 48:117–122.

9. Garcia-Armisen T, Papalexandratou Z, Hendryckx H, Camu N, Vrancken G, De Vuyst L, Cornelis P. 2010. Diversity of the total bacterial community associated with Ghanaian and Brazilian cocoa bean fermentation samples as revealed by a 16 S rRNA gene clone library. Appl Microbiol Biotechnol 87:2281–2292.

10. Blaxter M, Mann J, Chapman T, Thomas F, Whitton C, Floyd R, Abebe E. 2005. Defining operational taxonomic units using DNA barcode data. Philos Trans R Soc B Biol Sci 360:1935–1943.

11. Schloss PD, Westcott SL, Ryabin T, Hall JR, Hartmann M, Hollister EB, Lesniewski RA, Oakley BB, Parks DH, Robinson CJ, Sahl JW, Stres B, Thallinger GG, Van Horn DJ, Weber CF. 2009. Introducing mothur: Open-source, platform-independent, community-supported software for describing and comparing microbial communities. Appl Environ Microbiol 75:7537–7541.

12. Caporaso JG, Kuczynski J, Stombaugh J, Bittinger K, Bushman FD, Costello EK, Fierer N, Peña AG, Goodrich JK, Gordon JI, Huttley G a, Kelley ST, Knights D, Koenig JE, Ley RE, Lozupone C a, Mcdonald D, Muegge BD, Pirrung M, Reeder J, Sevinsky JR, Turnbaugh PJ, Walters W a, Widmann J, Yatsunenko T, Zaneveld J, Knight R. 2010. Correspondence QIIME allows analysis of high-throughput community sequencing data Intensity normalization improves color calling in SOLiD sequencing. Nat Publ Gr 7:335– 336.

13. Eren AM, Maignien L, Sul WJ, Murphy LG, Grim SL, Morrison HG, Sogin ML. 2013. Oligotyping: Differentiating between closely related microbial taxa using 16S rRNA gene data. Methods Ecol Evol 4:1111–1119.

14. Eren AM, Borisy GG, Huse SM, Mark Welch JL. 2014. Oligotyping analysis of the human oral microbiome. Proc Natl Acad Sci 111:E2875–E2884.

15. Maignien L, DeForce EA, Chafee ME, Murat Eren A, Simmons SL. 2014. Ecological succession and stochastic variation in the assembly of *Arabidopsis thaliana* phyllosphere communities. MBio 5:1–10.

16. Canani RB, Sangwan N, Stefka AT, Nocerino R, Paparo L, Aitoro R, Calignano A, Khan AA, Gilbert JA, Nagler CR. 2016. Lactobacillus rhamnosus GG-supplemented formula expands butyrate-producing bacterial strains in food allergic infants. ISME J 10:742–750.

17. De Filippis F, Troise AD, Vitaglione P, Ercolini D. 2018. Different temperatures select distinctive acetic acid bacteria species and promotes organic acids production during Kombucha tea fermentation. Food Microbiol 73:11–16.

18. Wu L, Wen C, Qin Y, Yin H, Tu Q, Van Nostrand JD, Yuan T, Yuan M, Deng Y, Zhou J. 2015. Phasing amplicon sequencing on Illumina Miseq for robust environmental microbial community analysis. BMC Microbiol 15:125.

19. Evaluaciones Agropecuarias Municipales -EVA-Oficina de planeación y prospectiva MADR. 2016. Evaluaciones Agropecuarias Municipales: cultivo de Cacao - año 2016. Evaluaciones Agropecu del Minist Agric y Desarro Rural.

20. Federación Nacional de Cacaoteros. 2013. Guía ambiental para el cultivo del cacao 1–126.

21. Magalhães da Veiga Moreira I, de Figueiredo Vilela L, da Cruz Pedroso Miguel MG, Santos C, Lima N, Freitas Schwan R. 2017. Impact of a Microbial Cocktail Used as a Starter Culture on Cocoa Fermentation and Chocolate Flavor. Molecules 22: 1–15.

22. Faith JJ, Guruge JL, Charbonneau M, Subramanian S, Seedorf H, Goodman AL, Clemente JC, Knight R, Heath AC, Leibel RL, Rosenbaum M, Gordon JI. 2013. The long-term stability of the human gut microbiota. Science 341:1–19.

23. Caporaso JG, Lauber CL, Walters WA, Berg-Lyons D, Lozupone CA, Turnbaugh PJ, Fierer N, Knight R. 2011. Global patterns of 16S rRNA diversity at a depth of millions of sequences per sample. Proc Natl Acad Sci U S A 108 Suppl:4516–22.

24. Toju H, Tanabe AS, Yamamoto S, Sato H. 2012. High-coverage ITS primers for the DNA-based identification of ascomycetes and basidiomycetes in environmental samples. PLoS One 7:1–11.

25. Caro-Quintero A, Ochman H. 2015. Assessing the unseen bacterial diversity in microbial communities. Genome Biol Evol 7:3416–3425.

26. Knights D, Kuczynski J, Charlson ES, Zaneveld J, Mozer MC, Collman RG, Bushman FD, Knight R, Kelley ST. 2011. Bayesian community-wide culture-independent microbial source tracking. Nat Methods 8:761–765.

27. Sokal RR, Oden NL. 1978. Spatial autocorrelation in biology: 1. Methodology. Biol J Linn Soc 10:199–228.

28. Hammer Ø, Harper DAT, Ryan PD. 2001. PAST: paleontological statistics software package for education and data analysis. Palaeontol Electron 4:9.

29. Papalexandratou Z, Vrancken G, De Bruyne K, Vandamme P, De Vuyst L. 2011. Spontaneous organic cocoa bean box fermentations in Brazil are characterized by a restricted species diversity of lactic acid bacteria and acetic acid bacteria. Food Microbiol 28:1326–1338.

30. Pereira GV de M, Soccol VT, Soccol CR. 2016. Current state of research on cocoa and coffee fermentations. Curr Opin Food Sci 7:50–57.

31. Papalexandratou Z, Falony G, Romanens E, Jimenez JC, Amores F, Daniel HM, De Vuyst L. 2011. Species diversity, community dynamics, and metabolite kinetics of the microbiota associated with traditional ecuadorian spontaneous cocoa bean fermentations. Appl Environ Microbiol 77:7698–7714.

32. Ardhana MM, Fleet GH. 2003. The microbial ecology of cocoa bean fermentations in Indonesia. Int J Food Microbiol 86:87–99.

33. Papalexandratou Z, Lefeber T, Bahrim B, Lee OS, Daniel HM, De Vuyst L. 2013. *Hanseniaspora opuntiae*, *Saccharomyces cerevisiae*, *Lactobacillus fermentum*, and *Acetobacter pasteurianus* predominate during well-performed Malaysian cocoa bean box fermentations, underlining the importance of these microbial species for a successful cocoa. Food Microbiol 35:73–85.

34. Arana-Sánchez A, Segura-García LE, Kirchmayr M, Orozco-Ávila I, Lugo-Cervantes E, Gschaedler-Mathis A. 2015. Identification of predominant yeasts associated with artisan Mexican cocoa fermentations using culture-dependent and culture-independent approaches. World J Microbiol Biotechnol 31:359–369.

35. Cocolin L, Alessandria V, Dolci P, Gorra R, Rantsiou K. 2013. Culture independent methods to assess the diversity and dynamics of microbiota during food fermentation. Int J Food Microbiol 167:29–43.

36. Nielsen DS, Teniola OD, Ban-Koffi L, Owusu M, Andersson TS, Holzapfel WH. 2007. The microbiology of Ghanaian cocoa fermentations analysed using culture-dependent and culture-independent methods. Int J Food Microbiol 114:168–186.

37. Schwan RF, Wheals AE. 2004. The microbiology of cocoa fermentation and its role in chocolate quality. Crit Rev Food Sci Nutr 44:205–221.

38. Reuter G. 1985. Elective and selective media for lactic acid bacteria. Int J Food Microbiol 2:55–68.

39. Eren AM, Morrison HG, Lescault PJ, Reveillaud J, Vineis JH, Sogin ML. 2015. Minimum entropy decomposition: Unsupervised oligotyping for sensitive partitioning of high-throughput marker gene sequences. ISME J 9:968–979.

40. Adler P, Bolten CJ, Dohnt K, Hansen CE, Wittmann C. 2013. Core fluxome and metafluxome of lactic acid bacteria under simulated cocoa pulp fermentation conditions. Appl Environ Microbiol 79:5670–5681.

41. Papalexandratou Z, Camu N, Falony G, De Vuyst L. 2011. Comparison of the bacterial species diversity of spontaneous cocoa bean fermentations carried out at selected farms in Ivory Coast and Brazil. Food Microbiol 28:964–973.

42. Pereira GVDM, Magalhães-Guedes KT, Schwan RF. 2013. RDNA-based DGGE analysis and electron microscopic observation of cocoa beans to monitor microbial diversity and distribution during the fermentation process. Food Res Int 53:482–486.

43. Schwan RF. 1998. Cocoa fermentations conducted with a defined microbial cocktail inoculum. Appl Environ Microbiol 64:1477–1483.

44. Batista NN, Ramos CL, Dias DR, Pinheiro ACM, Schwan RF. 2016. The impact of yeast starter cultures on the microbial communities and volatile compounds in cocoa fermentation and the resulting sensory attributes of chocolate. J Food Sci Technol 53:1101–1110.

